# Transfer of the synechan biosynthesis and regulatory pathway enables sulfated polysaccharide production in *Synechococcus elongatus* PCC 7942

**DOI:** 10.64898/2025.12.18.694786

**Authors:** Kaisei Maeda, Kazuma Ohdate, Yutaka Sakamaki, Kaori Nimura-Matsune, Satoru Watanabe

**Author notes:** **Correspondence**: Co-corresponding Author Kaisei Maeda Satoru Watanabe.

## Abstract

Many sulfated polysaccharides (SPSs) possess useful physicochemical properties and biological activities and are widely used in industry. Currently, major SPSs are derived from livestock and marine organisms. To contribute both to addressing global environmental challenges and to the development of diverse SPSs, microbial production systems are therefore required. Nevertheless, bacteria capable of producing SPSs are limited. In contrast, many cyanobacteria synthesize diverse SPSs, making them promising hosts to produce valuable SPSs, sources for novel SPSs, and genetic resources for SPS biosynthesis. However, no study has introduced an entire heterologous SPS biosynthetic system into cyanobacteria to produce a heterologous SPS. We recently identified a novel SPS, synechan, and elucidated the genetic basis for the biosynthesis and regulation of cyanobacterial SPSs. Here, we heterologously expressed the gene set responsible for synechan biosynthesis and its regulation in the non-SPS-producing cyanobacterium *Synechococcus elongatus* PCC 7942 and successfully induced the production of an SPS. This work provides a foundation for engineering cyanobacteria to produce useful SPSs and, to our knowledge, the first functional heterologous reconstitution in cyanobacteria of a large gene cluster encoding a complex membrane-associated biosynthetic system and thus an important step in synthetic-biology-based engineering of cyanobacteria.

## Introduction

Sulfated polysaccharides (SPSs), polysaccharides modified with sulfate groups, exhibit favorable physicochemical properties such as high water-retention capacity and viscosity, as well as various biological activities including anti-inflammatory and antiviral effects^1^. Among them, the animal-derived glycosaminoglycans chondroitin sulfate and heparin are well known and are mainly used as ingredients in cosmetics and pharmaceuticals. Thus, SPSs represent particularly high-value members of the polysaccharide family. At present, most industrially used SPSs are obtained from livestock and aquatic animals such as sharks, as well as from seaweeds. From the perspectives of addressing global environmental challenges, transitioning toward nature-positive production systems and the development of diverse SPSs, alternative SPS production platforms are being sought.

When the carbohydrate backbone of a desired SPS is available and the corresponding specific sulfotransferases are known, in vitro sulfation is in principle possible. However, the sulfate donor 3’-phosphoadenosine-5’-phosphosulfate (PAPS), which is required as the substrate for sulfotransferases, is expensive, making such in vitro processes impractical at an industrial scale. Consequently, efforts have focused on producing chondroitin sulfate and related SPSs in vivo using genetically engineered microorganisms^2^. Chondroitin sulfate is not only a valuable polysaccharide, but is also highly amenable to bacterial genetic engineering because *Escherichia coli* K-4 naturally synthesizes a polysaccharide backbone that is almost identical to the chondroitin backbone^3^. In representative studies, host bacteria such as *E. coli* have been engineered to express the *E. coli* K-4 chondroitin backbone biosynthetic gene cluster together with optimized animal-derived sulfotransferases to produce chondroitin sulfate^4^. However, these systems still suffer from low degrees of sulfation and low productivity and have not yet been implemented industrially. One possible reason is that the host microorganisms used in these studies do not naturally produce SPSs. Although many bacteria synthesize acidic polysaccharides, bacterial species that produce SPSs appear to be restricted to cyanobacteria and a limited number of marine bacteria^5,6^. As a result, compared with other bacterial acidic polysaccharides such as xanthan, our knowledge of the biosynthesis and regulation of bacterial SPSs remains very limited.

Cyanobacteria are oxygenic photosynthetic prokaryotes that inhabit diverse environments and play important roles in global ecosystems ^7,8^. In recent years, they have also attracted attention as chassis organisms for photosynthesis-driven bioproduction^9,10^. In nature, cyanobacteria form a variety of multicellular structures and biofilms, and diverse SPSs are known as major components of these extracellular matrices^11^. Major examples of cyanobacterial sulfated polysaccharides are spirulan from *Arthrospira platensis* (vernacular name, “Spirulina”), sacran from *Aphanothece sacrum* (vernacular name, “Suizenji-Nori”) and cyanoflan from *Cyanothece* sp. CCY 0110 ^12–14^. Sacran exhibits high water-retention capacity and anti-inflammatory activity, and has already been implemented industrially, for example as a cosmetic ingredient^15^. However, although the diversity and utility of cyanobacterial SPSs have been recognized and some have been developed for applications, very little is known about the mechanisms underlying SPS biosynthesis and regulation, and only a few genetic engineering approaches have been applied to SPS production. We have long been investigating the biosynthesis and regulatory mechanisms of cyanobacterial extracellular polysaccharides^16^. Recently, we discovered that the freshwater model cyanobacterium *Synechocystis* sp. PCC 6803 (*S.*6803) accumulates viscous exopolysaccharides (EPSs) and forms bloom-like cell aggregates, and we used this phenomenon as a clue to comprehensively identify a novel SPS, synechan, together with its biosynthetic and regulatory gene cluster, designated *xss*^6^. The *xss* gene cluster represents the first elucidated cyanobacterial SPS biosynthetic system. Database analyses of cyanobacterial genome sequences based on the *xss* information suggested that cyanobacterial SPS biosynthetic systems generally consist of combinations of canonical bacterial exopolysaccharide biosynthetic machineries, Wzx/Wzy-dependent or ABC transporter-dependent pathways, and sulfotransferases. This finding implies that, in principle, it should be possible to engineer cyanobacteria to produce a variety of SPSs by combining bacterial polysaccharide biosynthetic pathways with appropriate sulfotransferases. However, to our knowledge, no such metabolic engineering studies in cyanobacteria have yet been reported. Moreover, genome analyses have revealed numerous gene clusters on cyanobacterial genomes that are predicted to encode diverse SPS biosynthetic systems, but most of the corresponding SPS-producing cyanobacterial species are non-model organisms. Therefore, omics-based or synthetic biology approaches will be essential to elucidate the functions of these genes.

Given this background, functionally expressing known useful SPS biosynthetic systems, candidate SPS biosynthetic gene clusters, or rationally designed SPS pathways in cyanobacteria is crucial both for basic research aimed at understanding the biosynthetic mechanisms and functions of cyanobacterial SPSs and for applied research aimed at producing valuable SPSs. In this study, as a first step toward this goal, we examined whether the cyanobacterial SPS biosynthetic system Xss that we previously identified in *S.*6803 can be functionally expressed in another model cyanobacterium, *Synechococcus elongatus* PCC 7942 (*S*.7942) (Fig. 1). To our knowledge, there has been no previous report in cyanobacteria of an entire large gene cluster encoding such a complex membrane-associated biosynthetic system being heterologously expressed and functionally reconstituted.

**Figure 1.**
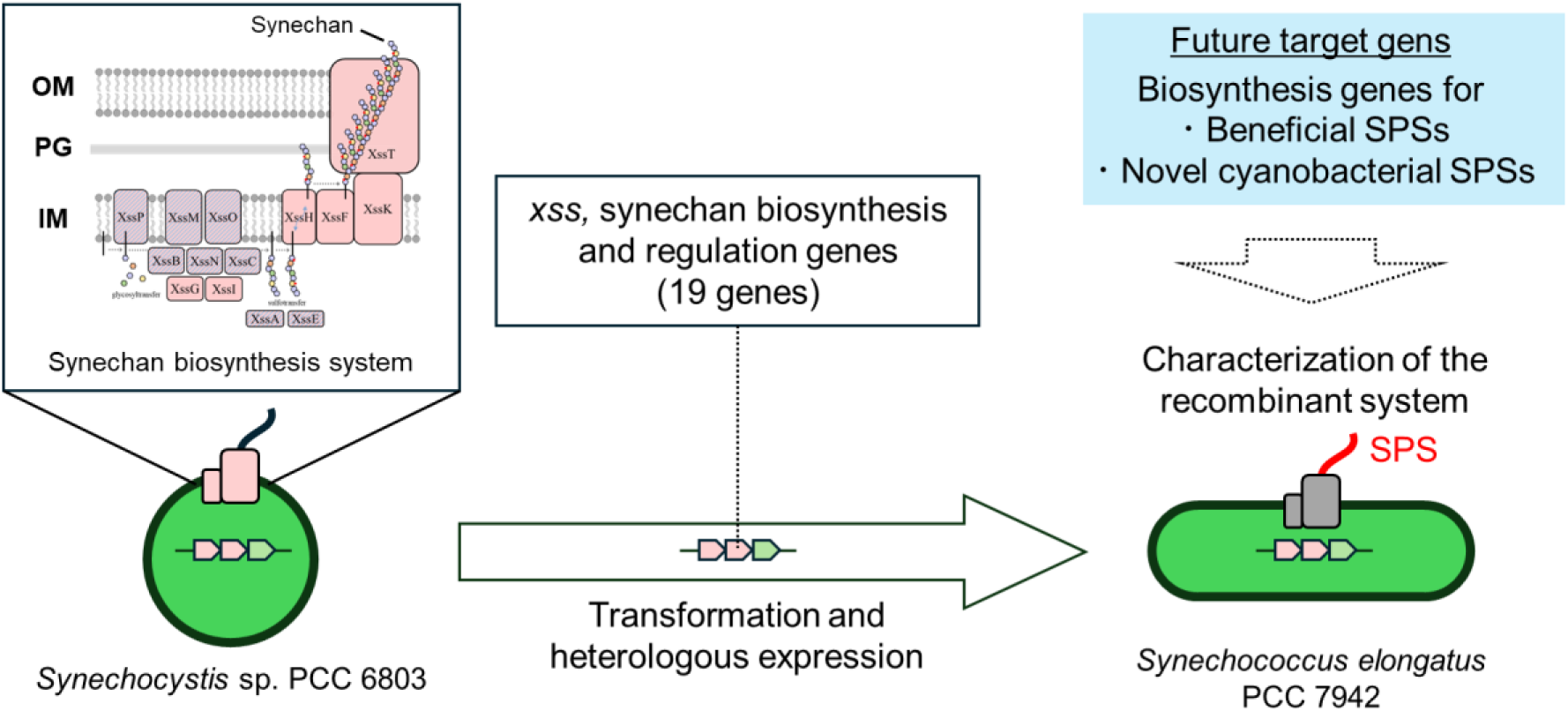
Graphical abstract of this study. We heterologously expressed the synechan biosynthetic genes from *Synechocystis* sp. PCC 6803 (*S.*6803), together with associated regulatory genes, in *Synechococcus elongatus* PCC 7942 (*S*.7942), a model cyanobacterium that does not natively produce sulfated polysaccharides, thereby enabling sulfated polysaccharide production. This result demonstrates the feasibility of functionally reconstituting biosynthetic gene sets for the heterologous production of valuable or novel sulfated polysaccharides in cyanobacteria. OM, outer membrane; PG, peptidoglycan; IM, inner membrane; SPS, sulfated polysaccharide.

## Results

### Design of heterologous expression of the synechan biosynthetic *xss* gene cluster

The synechan biosynthetic and regulatory gene cluster *xss* in *S.*6803 consists of the biosynthetic genes *xssA-xssP* and *xssT*, which are directly involved in synechan production, and the regulatory genes *xssQ*, *xssR*, and *xssS*, which control their expression^6^. The genes *xssA-xssS* are clustered on the megaplasmid pSYSM in *S.*6803, but they do not form a single operon^17^ (Fig. S1). In contrast, *xssT* is located on the chromosome and is thought to function also in other chromosomally encoded capsular polysaccharide (CPS) biosynthetic systems. Previous studies have suggested that, in the regulatory system consisting of the sensor histidine kinase XssQ, the response regulator XssR, which lacks a DNA-binding domain, and the cyanobacteria-specific transcriptional regulator XssS, XssQ acts as the direct transcriptional regulator of synechan biosynthesis, XssR positively regulates XssQ activity, and XssS suppresses XssQ-dependent transcription via XssR^6^. Among the *xss* genes, only the genes encoding glycosyltransferases and sulfotransferases are directly regulated by XssQ (Fig. S1), and this regulation leads to marked changes in synechan production.

Based on these characteristics of the *xss* gene cluster, we designed a strategy to heterologously express the synechan biosynthetic system in *S*.7942. To express *xssA-xssP* as a single operon under a strong promoter, it would be necessary to align the orientation of all genes and insert appropriate ribosome-binding sites (RBSs) between them, which would require considerable effort. Therefore, we adopted an alternative strategy in which the *trc* promoter was placed upstream of *xssP*, encoding the enzyme that catalyzes the initial glycosyl transfer reaction in synechan biosynthesis, and the expression of the remaining genes was expected to be induced by heterologous expression of *xssQ* and *xssR*. Specifically, we constructed pYS1C-*xssP-A* by cloning *xssA-xssP* into our previously developed high-expression shuttle vector pYS1C^18^, and pBNS1-*xssQRT* by cloning *xssQ*, *xssR*, and *xssT*, linked in an operon via synthetic RBSs, into the neutral site 1 (NS1) locus of the chromosome (Fig. 2a and S2). In pYS1C-*xssP-A*, *xssP* is placed under the control of the *trc* promoter, and in NS1-xssQRT, the xssQ-xssR-xssT operon is also under the control of the *trc* promoter; thus, both cassettes are transcriptionally induced by IPTG.

**Figure 2.**
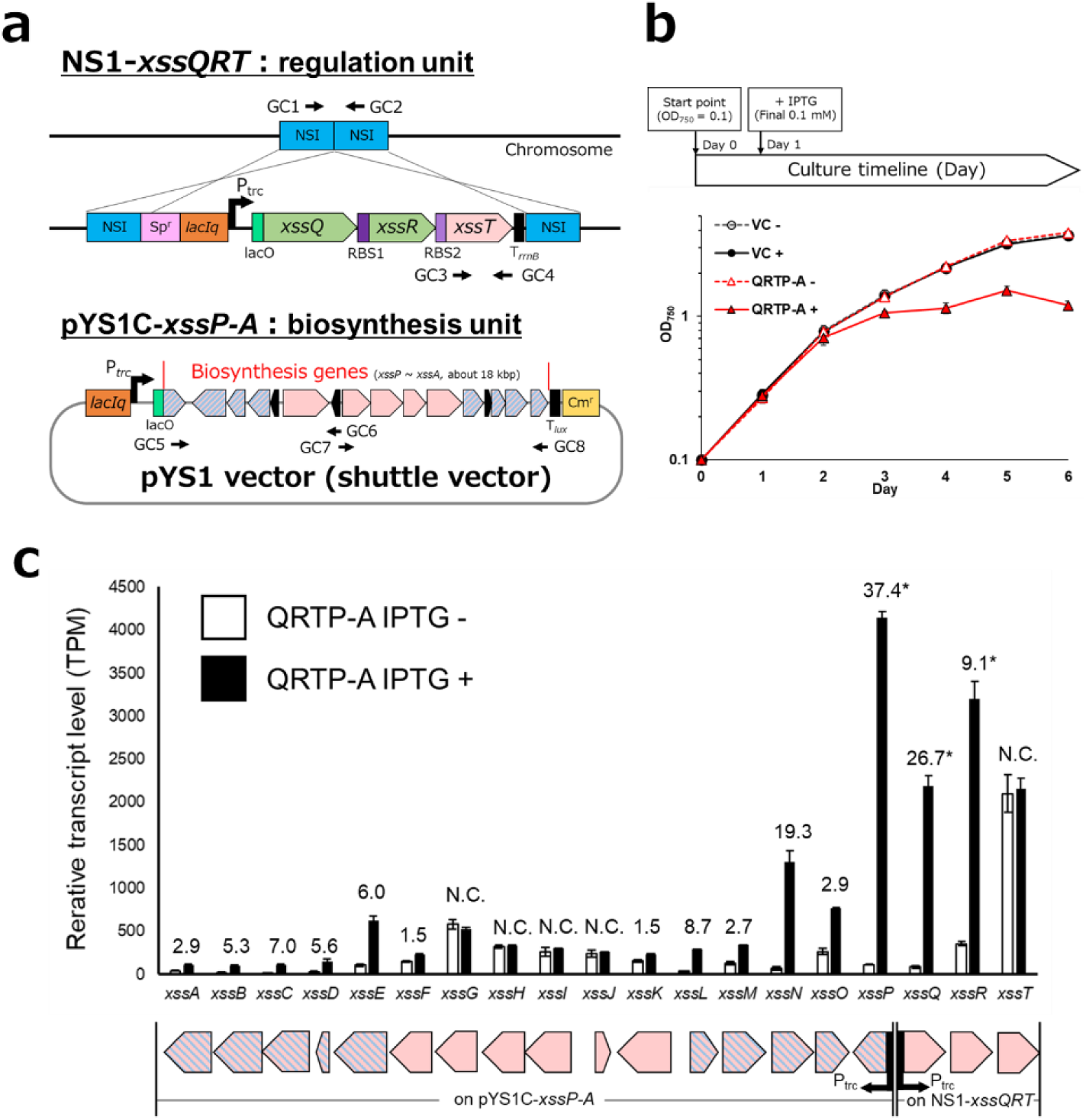
Construction and gene expression analysis of S.7942 mutants carrying transferred *xss* genes. (a) Schematic representation of the strategy used for heterologous expression of the synechan biosynthetic xss gene cluster. The upper panel shows the introduction of the transcriptional regulatory genes *xssQ* and *xssR* and *xssT*, encoding the outer-membrane polysaccharide export protein (OPX) required for synechan secretion, into the chromosomal neutral site NS1 using pBNS1-*xssQRT*. The lower panel shows the introduction of the synechan biosynthetic genes *xssA*-*xssP* using the shuttle vector pYS1C-*xssP-A*. GC1-8 indicate the primer sets used for genotyping and segregation analysis of the resultant mutant strains (Table S1). (b) Growth curves of the *S*.7942 strain transformed with pYSC1-GFP (VC) and the heterologous synechan biosynthetic expression strain (QRTP-A) in the presence or absence of IPTG. (c) Gene expression analysis of the xss gene cluster based on RNA-seq of cells harvested 2 days after IPTG induction. White bars and black bars represent transcript levels without and with IPTG induction, respectively. Labels above each gene indicate either no significant change between the IPTG conditions (N.C.) or the fold change. Labels marked with an asterisk indicate data for genes that are under the control of the Ptrc promoter. The schematic below the graph shows the arrangement of the *xss* genes corresponding to the bars in the graph. Each pentagon represents a gene; pentagons with pink and blue stripes indicate genes that are under XssQ-dependent transcriptional regulation in *S.*6803, and pink pentagons indicate the remaining *xss* genes. Error bars indicate SD (n = 3).

To construct the engineered strains, we first introduced the pBNS1-*xssQRT* construct into the wild-type *S*.7942 by natural transformation. The resulting strain is hereafter referred to as QRT. Using QRT as the parental strain, we then introduced the pYS1C-*xssP-A* construct to obtain the QRTP-A strain. The genotypes of these strains and complete segregation of the chromosomal copies were confirmed by PCR amplification of specific genomic regions followed by agarose gel electrophoresis (Table S1 and Fig. S3). As a result, at least three independent, fully segregated transformants were obtained for each strain.

### Expression of introduced *xss* genes in the QRTP-A strain

To examine the expression of the introduced *xss* genes in the constructed QRTP-A strain, we extracted total RNA from cells 2 days after IPTG induction (Fig. 2b) and performed RNA-seq analysis (Fig. 2c). For the genes integrated into the NS1 locus on the chromosome, *xssQ* and *xssR* showed strong transcriptional induction under the IPTG-induced condition, as expected. In contrast, *xssT* exhibited high transcript levels regardless of IPTG addition, suggesting the presence of an unknown strong promoter in the upstream region of *xssR*. For the genes located on the shuttle vector pYS1C-*xssP-A*, those that are not under XssQ-dependent transcriptional control in *S.*6803 (indicated in pink) maintained similar transcript levels irrespective of IPTG addition. By contrast, the genes that are under XssQ-dependent control in *S.*6803 (*xssA*, *xssB*, *xssC*, *xssD*, *xssE*, *xssL*, *xssM*, *xssN*, and *xssO*) showed significantly increased transcript levels under the IPTG-induced condition. These results indicate that XssQ/XssR-dependent transcriptional regulation of *xss* genes is functional in the heterologous host *S.*7942.

The triplet of regulatory genes *xssQ*, *xssR*, and *xssS* is widely conserved among SPS-producing cyanobacteria, and homologs of *xssQ* are also found in other cyanobacteria, although none are present in *S*.7942. In *S.*6803, it was previously unclear whether XssQ and XssR alone are necessary and sufficient for transcriptional regulation of the *xss* genes. Our results demonstrate that, at least within cyanobacteria, the presence of XssQ and XssR is sufficient for XssQ to bind consensus sequences upstream of its target genes and regulate their transcription. This implies that, in heterologous expression of SPS biosynthetic systems predicted to employ an XssS-XssR-XssQ-type regulatory module, co-expression of homologs of *xssQ* (and *xssR*) can be expected to enhance transcript levels of the corresponding target genes. Because Wzx/Wzy-type and ABC transporter-type polysaccharide biosynthetic systems typically comprise many genes, and in cyanobacterial genomes such genes are often dispersed across the genome rather than forming a single cluster^16,19^, there are substantial barriers to expressing them as contiguous gene clusters in heterologous hosts. A system that allows such gene sets to be introduced and expressed in a heterologous cyanobacterium while preserving their native genomic organization will therefore be highly useful for future research and development.

### Growth and EPS production in the QRTP-A strain

The QRTP-A strain and the previously established vector control strain VC^18^ were cultivated in liquid medium with or without IPTG to examine cell growth (Fig. 2b). In the VC strain, no significant difference in growth rate was observed between cultures with and without IPTG. In contrast, in the QRTP-A strain, growth under the non-induced condition (without IPTG) was comparable to that of the VC strain, whereas under the IPTG-induced condition, growth almost completely ceased within 1 day after IPTG addition, and the OD₇₅₀ value after 6 days was markedly lower than that under the non-induced condition. In short, expression of the *xss* genes negatively affected cell growth.

Under IPTG-induced conditions, the QRTP-A strain formed small cell aggregates even in bubbling cultures (Fig. 3a). In *S.*6803, when cultures that have accumulated synechan are left standing, cells float to the surface owing to synechan and photosynthetically generated gas bubbles, forming bloom-like viscous aggregates at the air-liquid interface^6^. In contrast, when QRTP-A cultures were left standing, the cells sedimented to the bottom of the vessel and the culture remained non-viscous. To investigate the cause of this aggregation, we observed the cells by scanning electron microscopy (Fig. 3b). Whereas the VC strain displayed no characteristic surface structures, numerous fibrous structures and granular materials attached to them were observed on the cell surface and in the extracellular space of the QRTP-A strain. This result suggests that high-molecular-weight polysaccharides are specifically produced and accumulated extracellularly in the QRTP-A strain. In addition, QRTP-A cells were markedly elongated, a phenotype that is commonly observed in rod-shaped cyanobacteria under stress^20^. We infer that this cell elongation, together with the accumulation of extracellular polymers, contributed to the observed cell aggregation.

**Figure 3.**
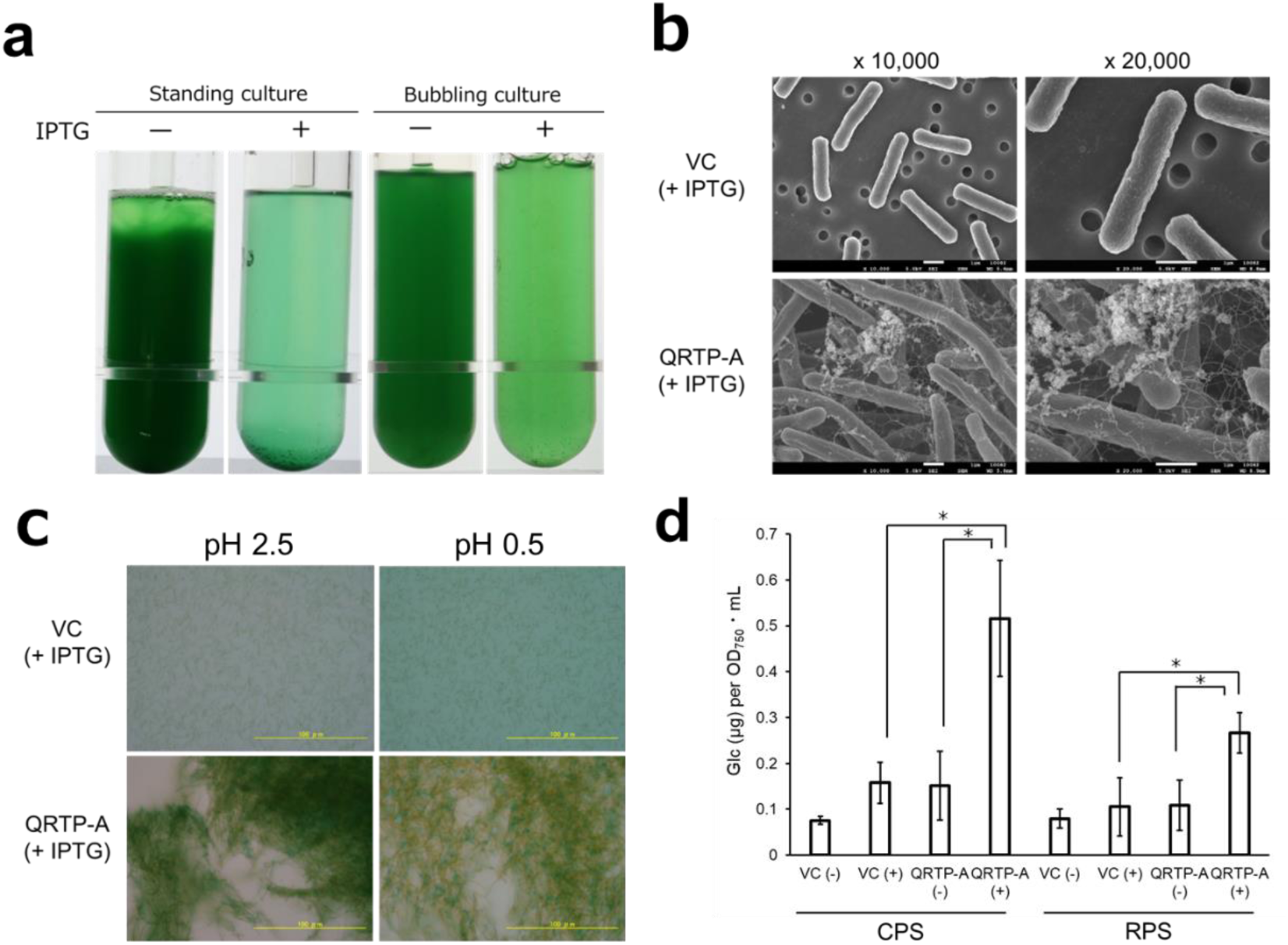
Phenotypic characterization of the *S.*7942 QRTP-A mutant strain. (a) Comparison of QRTP-A cultures with or without IPTG induction. “Standing culture” indicates cultures grown under the same conditions as in Fig. 2b for 6 days and then left standing for an additional 2 days, whereas “bubbling culture” indicates cultures grown with aeration and agitation under the same conditions for 3 days. (b) Scanning electron micrographs of the VC and QRTP-A strains 5 days after IPTG induction. Fibrous and aggregate-like structures were observed on the cell surface of the QRTP-A strain. (c) Alcian blue staining of the VC and QRTP-A strains 2 days after IPTG induction. Alcian blue stains acidic polysaccharides at pH 2.5 and specifically sulfated polysaccharides at pH 0.5. (d) Total carbohydrate content, expressed as glucose equivalents, of the cell-associated capsular polysaccharides (CPS) and released polysaccharides in the medium (RPS) fractionated from VC and QRTP-A strains 5 days after IPTG induction. Total sugars were quantified using the phenol-sulfuric acid method. n = 3 (error bars, SD). *, P < 0.05 (Welch’s t-test).

Next, to examine whether these fibrous structures consist of SPSs, we performed Alcian blue staining (Fig. 3c). In Alcian blue staining, acidic polysaccharides in general are stained blue at pH 2.5, whereas sulfated polysaccharides are selectively stained at pH 0.5. In QRTP-A cultures, blue aggregates were observed under both pH conditions. This result suggests accumulation of SPSs in the extracellular space. Notably, in the Alcian blue images, the distribution of stained SPSs appears more localized than the distribution of cells in the SEM images. This localization is considered to be an artifact caused by Alcian-blue-induced aggregation of acidic polysaccharides; in reality, the polymers are expected to be more loosely distributed around the cells^21^.

To quantitatively evaluate this polysaccharide accumulation, we measured the total carbohydrate content by the phenol-sulfuric acid method (Fig. 3d). Extracellular polysaccharides were fractionated into released polysaccharides in the medium (RPS) and cell-associated capsular polysaccharides (CPS), and their amounts were compared between VC and QRTP-A strains under IPTG-induced and non-induced conditions. Under IPTG-induced conditions, the QRTP-A strain showed a significant increase in the accumulation of both RPS and CPS, whereas the VC strain showed no such increase. Thus, introduction of the *xss* gene cluster resulted in a significant enhancement of extracellular polysaccharide production. However, even under conditions in which XssQ expression was induced, the productivity was comparable to the amount of synechan accumulated by the *S*.6803 wild-type strain under non-inducing conditions for XssQ.

### Characterization of EPS produced by the QRTP-A strain

To qualitatively characterize the EPS produced by the QRTP-A strain, we analyzed the monosaccharide composition and the degree of sulfation of the RPS fraction (Table 1). The heterologous polysaccharide had an approximate monosaccharide composition of Glc:Gal:Man = 1:2:1 and a degree of sulfation of 22.6%. This composition suggests that the polymer may be an SPS consisting of repeating units carrying, on average, one sulfate group per four monosaccharide residues. By contrast, the original synechan produced by *S.*6803 showed a monosaccharide composition of Glc:Gal:Man:Xyl = 5:1:1:1 and a degree of sulfation of 26.6%. Synechan is estimated to contain two sulfate groups per repeating unit of eight monosaccharides. Thus, the polysaccharide produced by the *S*.7942 QRTP-A strain had a degree of sulfation comparable to that of synechan and a similar set of constituent monosaccharides, but the relative monosaccharide ratios differed. In previous work on *S.*6803, among the individual knockouts of the eight glycosyltransferase genes, only the *xssP*, *xssB*, *xssM*, and *xssN* mutants showed a marked reduction in synechan production and loss of cell aggregation^6^. In addition, of the two sulfotransferase genes (*xssA* and *xssE*), complete loss of synechan production was observed only in the *xssA* mutant. These results suggest that some components of the synechan sugar chain are essential for polysaccharide synthesis, whereas others are dispensable and can be omitted without completely abolishing chain formation. In the QRTP-A strain analyzed in this study, the functions of the introduced *xss* gene set were likely not fully reproduced, and we infer that an incomplete sugar chain was synthesized. The facts that synechan is normally produced as an RPS whereas the QRTP-A polysaccharide accumulated mainly in the CPS fraction, and that its productivity was relatively low, may also reflect differences in sugar-chain composition from that of synechan.

**Table 1.** Chemical composition of RPS produced by the QRTP-A strain compared with synechan.

|  | *Sugars* |  |  |  |  |  |  |  |  |  |  | *Sulfate residues* |
| --- | --- | --- | --- | --- | --- | --- | --- | --- | --- | --- | --- | --- |
|  | Neutral sugars (mol/mol %) |  |  |  |  |  |  |  | Uronic acids (mol/mol %) |  |  | Substitution degree |
|  |  |  |  |  |  |  |  |  | Galacturonic acid | Glucuronic acid | Total | (mol/mol %) |
|  | Rhamnose | Ribose | Mannose | Fucose | Galactose | Xylose | Glucose | Total |  |  |  |  |
| *S. 7942* |  |  |  |  |  |  |  |  |  |  |  |  |
| QRTP-A | 4.2 | 2.1 | 23.5 | 2.5 | 44.3 | 0.1 | 23.4 | 100.0 | N.D. | N.D. | 0.0 | 22.5 |
| RPS |  |  |  |  |  |  |  |  |  |  |  |  |
| Synechan* | 13.1 | N.D. | 14.2 | 1.2 | 12.5 | 1.0 | 57.9 | 99.9 | N.D. | 0.1 | 0.1 | 26.6 |

N.D., not detected.
\* The data about synechan was taken in the previous research<sup>6</sup>

### Transcriptional changes caused by heterologous expression of the *xss* gene cluster

Based on the RNA-seq dataset, we analyzed transcriptional changes in the QRTP-A strain heterologously expressing the *xss* gene cluster. Using the thresholds │log₂FC│ ≥ 1 and FDR-adjusted *P* ≤ 0.05, 328 genes were significantly upregulated and 330 genes were significantly downregulated, indicating large-scale reprogramming of gene expression (Fig. 4, Tables 2, 3, S3 and S4). Among the sigma factors that play dominant roles in transcription, the primary σ factor *Synpcc7942_1557* (*rpoD1*) showed transcript levels reduced to approximately half of those in the control, whereas the group 2 σ factors *Synpcc7942_0672* (*rpoD3*) and *Synpcc7942_1746* (*rpoD2*) increased by about 4-fold and 2.1-fold, respectively. In addition, the transcript levels of *Synpcc7942_1849* (*rpoD5*), *Synpcc7942_1510* (*sigF1*), and *Synpcc7942_1784* (*sigF2*) were reduced to roughly half. These results suggest a shift in cellular metabolism from a RpoD1-dependent growth mode to a RpoD2/RpoD3-dominated stress-response mode^22^, which is consistent with the observation that cell growth was almost completely arrested under SPS-producing conditions. Moreover, disruption of *sigF* is known to induce biofilm formation in *S. elongatus* PCC 7942^23^ and to promote EPS accumulation in *Synechocystis* sp. PCC 6803^24^, and the QRTP-A strain is therefore likely to be in a similar physiological state.

**Figure 4.**
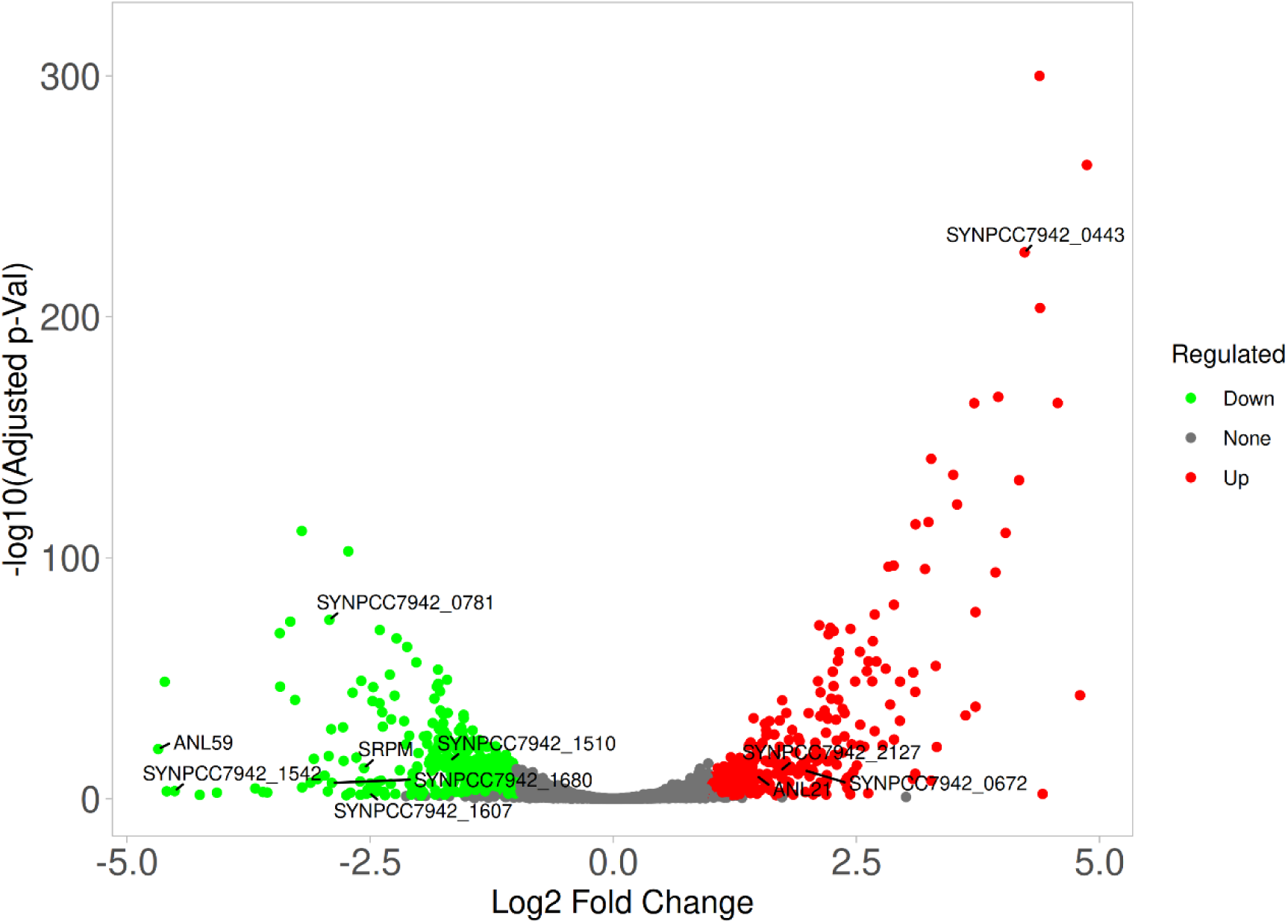
Volcano plot of transcriptomic changes in the QRTP-A strain. Upregulated and downregulated genes are depicted in red and green, respectively (│log₂FC│ ≥ 1, FDR-adjusted *P* ≤ 0.05). The x-axis shows log₂ fold change, and the y- axis shows −log₁₀ of the FDR-adjusted *P* value. Among the genes listed in Tables 2 and 3, those showing the largest changes in each category are indicated by labels.

**Table 2.** Representative genes significantly upregulated upon IPTG induction in the QRTP-A strain.

| Name | Product | Log2FC |
| --- | --- | --- |
| <u>Sigma factors</u> |  |  |
| Synpcc7942_0672 | RpoD3, group2 RNA polymerase sigma factor SigD | 1.98 |
| Synpcc7942_1746 | RpoD2, group2 RNA polymerase sigma factor SigB | 1.06 |
| <u>Membrane-associated protein coding genes</u> |  |  |
| Synpcc7942_0443 | conserved hypothetical protein | 4.25 |
| Synpcc7942_1870 | Secretion protein HlyD | 3.74 |
| Synpcc7942_1868 | conserved hypothetical protein | 3.73 |
| Synpcc7942_1869 | probable cation efflux system protein | 2.91 |
| Synpcc7942_0553 | Secretion protein HlyD | 2.71 |
| Synpcc7942_1904 | hemolysin secretion protein-like | 2.15 |
| Synpcc7942_1224 | ABC-transporter membrane fusion protein | 1.23 |
| Synpcc7942_1917 | permease of the drug/metabolite transporter | 1.01 |
| <u>Photosynthesis</u> |  |  |
| Synpcc7942_2127 | phycobilisome degradation protein NblA | 1.73 |
| Synpcc7942_1178 | photosystem II stability/assembly factor | 1.23 |
| Synpcc7942_1038 | photosystem II oxygen-evolving complex 23K protein | 1.12 |
| <u>pANL plasmid</u> |  |  |
| anL21 | COG3544; conserved hypothetical protein | 1.50 |

**Table 3.**
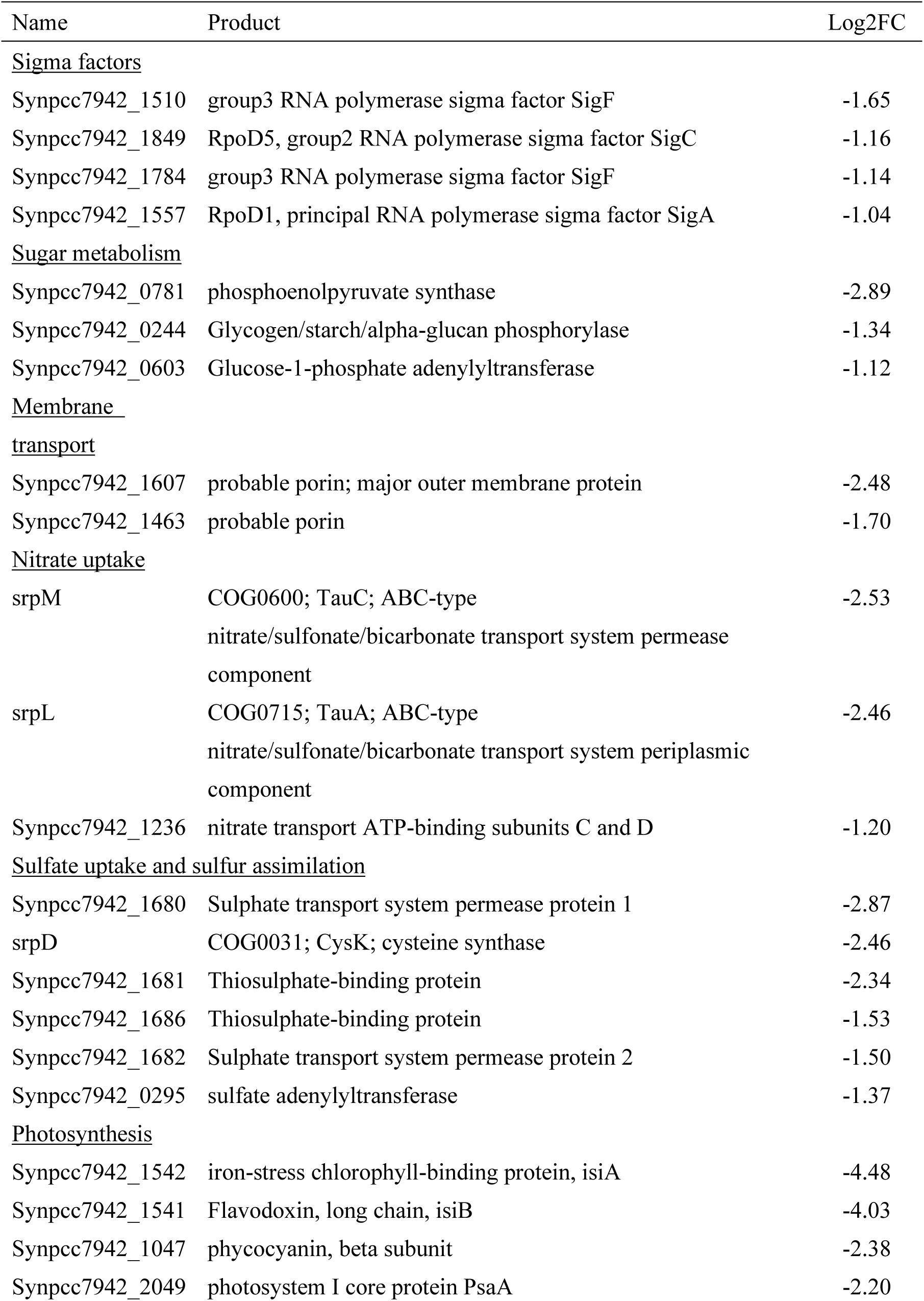
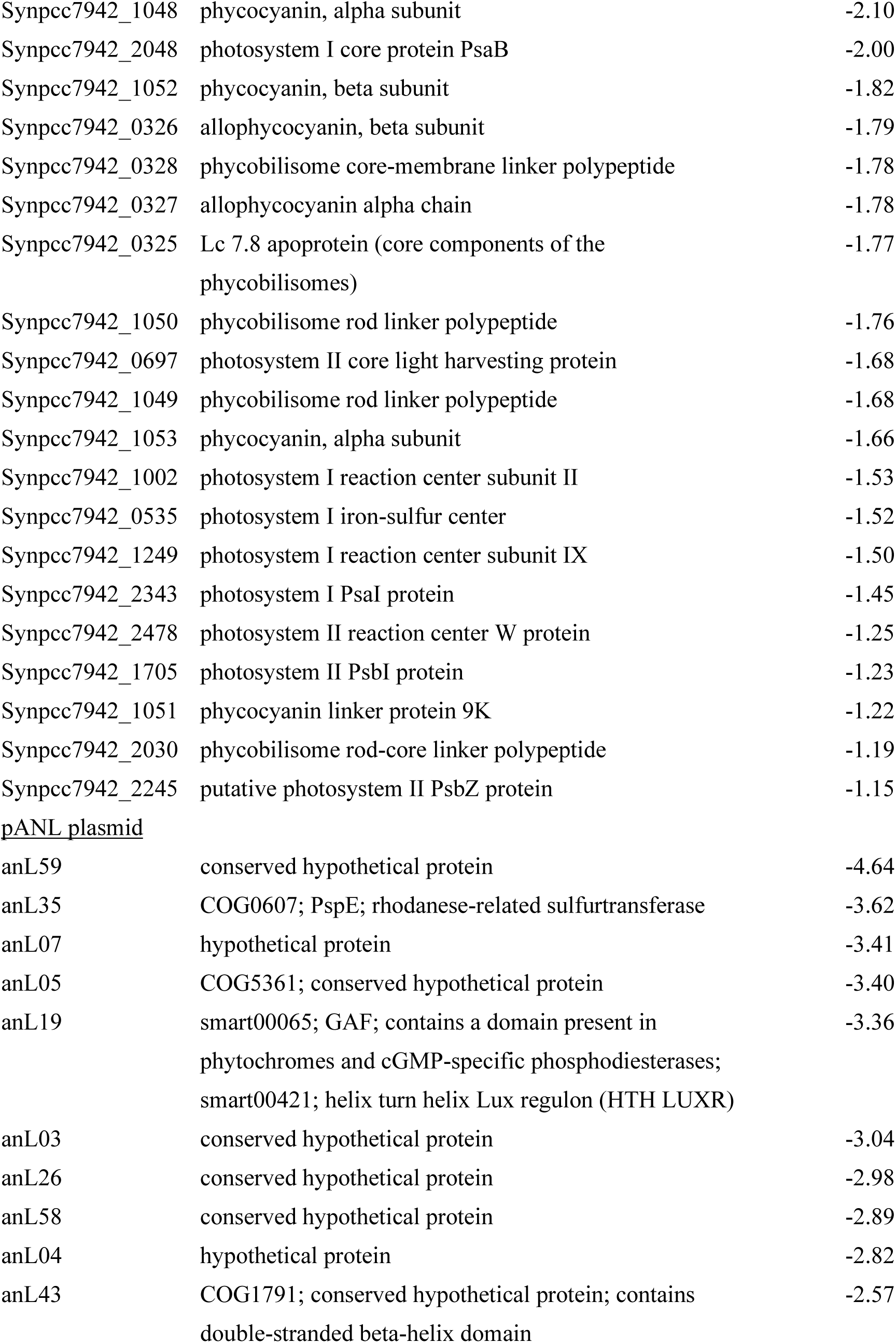
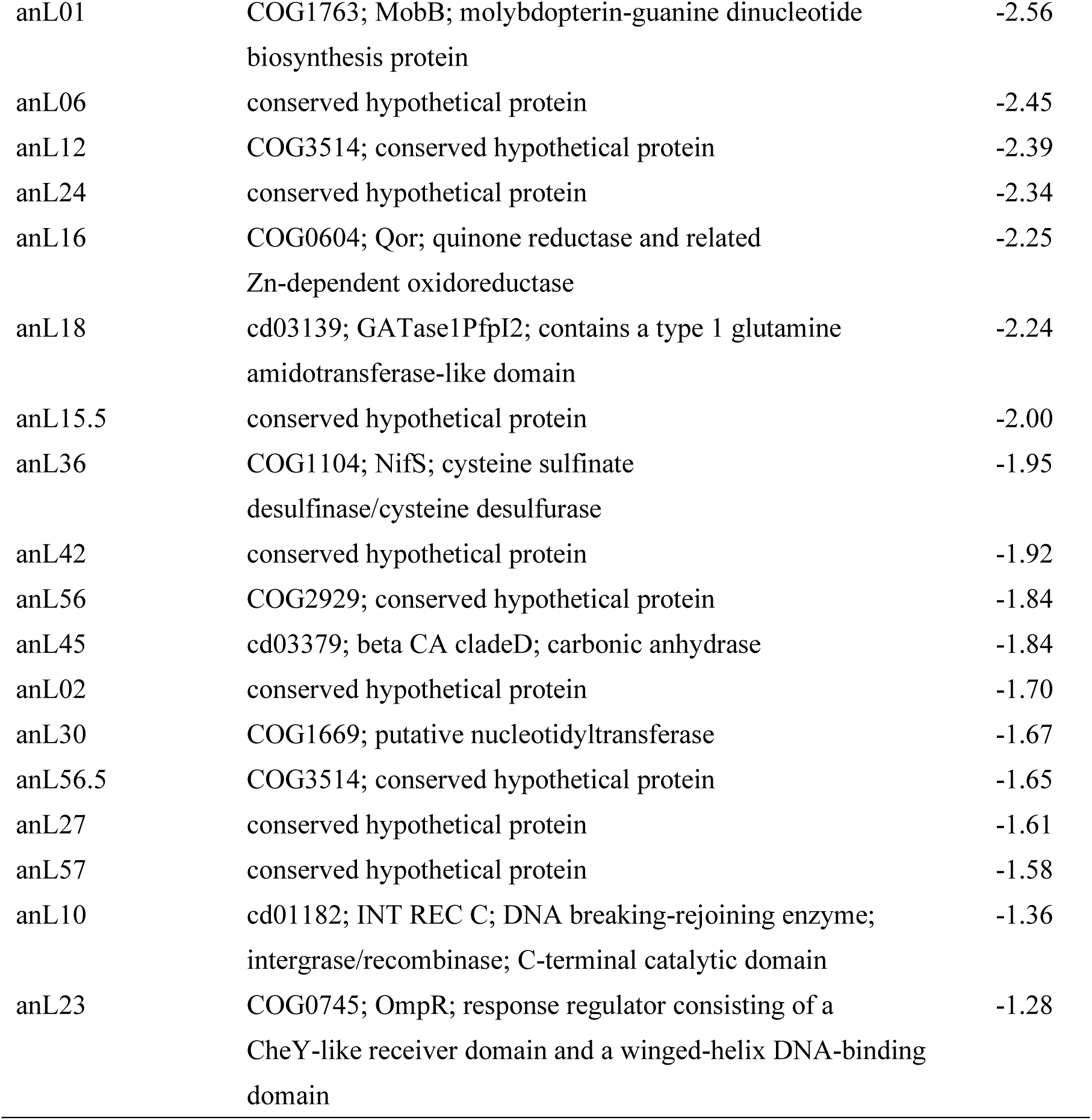
Representative genes significantly downregulated upon IPTG induction in the QRTP-A strain.

With respect to genes involved in polysaccharide synthesis and degradation, the transcript levels of the ADP-glucose pyrophosphorylase gene *Synpcc7942_0603*, which catalyzes the formation of the substrate for glycogen synthesis, and *Synpcc7942_0244*, which is involved in glycogen degradation, were both decreased (log₂FC = −1.12 and −1.34, respectively). In synechan, the constituent monosaccharides glucose and galactose are supplied as UDP-glucose and UDP-galactose, respectively, with UDP-galactose being produced by isomerization of UDP-glucose. UDP-glucose not only serves as a precursor for synechan biosynthesis but is also essential as a substrate for thylakoid membrane lipid synthesis in cyanobacteria^25^. Because UDP-glucose and ADP-glucose share glucose 1-phosphate as a precursor^26^, it is likely that flux toward ADP-glucose is downregulated to prioritize UDP-glucose synthesis. In addition, the transcript level of PEP (phosphoenolpyruvate) synthase *Synpcc7942_0781*, which generates PEP from pyruvate as a precursor for NDP-sugar synthesis and other reactions, was also markedly reduced (log₂FC = −2.9). This suggests the presence of metabolic regulation that prevents excessive allocation of carbon toward EPSs and storage polysaccharides. From the perspective of further improving heterologous EPS productivity, it will be necessary to engineer metabolism so that the sugars required for growth are secured while maintaining sufficient NDP-sugar supply.

Among membrane-related genes, we observed increased expression of several type I secretion systems and multidrug efflux pumps (*Synpcc7942_1868-1870*, *Synpcc7942_0553*, *Synpcc7942_1904*, *Synpcc7942_1224*, *Synpcc7942_1917*), along with decreased expression of porin-like genes (*Synpcc7942_1607*, *Synpcc7942_1463*). In addition, the transcript level of *Synpcc7942_0443*, which encodes a protein containing an SLH domain associated with the S-layer^27^, was strongly increased (log₂FC = 4.25). Because the Xss system is a biosynthetic system that includes many membrane proteins, these changes suggest that its heterologous expression substantially altered the structure and protein composition of the cell envelope.

Other membrane-associated genes also exhibited characteristic changes. The transcript levels of genes involved in nitrate uptake (*Synpcc7942_1236*, *srpM*, *srpL*)^28^ and those involved in sulfate uptake and sulfur assimilation (*Synpcc7942_1680*, *Synpcc7942_1681*, *Synpcc7942_1682*, *Synpcc7942_1686*, *srpD*, *Synpcc7942_0295*, and others) were decreased. These changes likely reflect the fact that the cells had largely exited an active growth state. From the standpoint of enhancing SPS production, however, it would be preferable to maintain high levels of sulfate uptake and supply of PAPS, the sulfate donor for sulfotransferases, and metabolic engineering to achieve this may be beneficial.

For photosynthesis-related genes, the transcript levels of phycobilisome components were broadly reduced, whereas transcription of *nblA* (*Synpcc7942_2127*), which promotes phycobilisome degradation^29^, was increased, indicating a reduction in antenna size. Many genes encoding components of photosystem I were downregulated. In photosystem II, the transcripts for CP43 and *psbI*, *psbW*, and *psbZ* were decreased, whereas *Synpcc7942_1178* (a PSII stability/assembly factor) and *Synpcc7942_1038* (PSII oxygen-evolving complex 23K protein) were upregulated. Taken together, these trends suggest that QRTP-A cells have largely shifted away from active photosynthesis. Furthermore, the iron-stress response genes *isiA* and *isiB*^30^ were strongly repressed under SPS-producing conditions (log₂FC ≈ −4.5 and −4.0, respectively), indicating that the cells were far from entering an iron-stress mode.

Another notable feature was the behavior of genes located on the endogenous plasmid pANL. Of the 39 pANL-encoded genes detected by RNA-seq, 29 showed significant expression changes, and among these, 28 genes were downregulated, with *anL21* being the only exception. We infer that these changes reflect competition for cellular resources required for plasmid maintenance, caused by introduction of the high-copy shuttle vector pYS1C-*xssP-A*. If this is the case, then in situations where genes on endogenous plasmids are important for host cyanobacterial growth or for the function of the introduced biosynthetic pathways, the use of high-copy shuttle vectors such as pYS1C may carry a risk.

## Discussion

Photosynthesis-driven biomanufacturing using algae and cyanobacteria has attracted considerable attention as a technology that can simultaneously achieve CO₂ fixation and the use of renewable energy. However, life cycle assessment and techno-economic analyses have repeatedly shown that fuel-only processes based on lipids or biofuels struggle to be economically viable because of the low unit value of these products^31,32^. A promising strategy to address this limitation is the “high-value co-product” or “algal biorefinery” concept, in which multiple high-value compounds are co-produced to help offset fuel-production costs^33^. Owing to their high water-retention capacity, viscosity and diverse biological activities, SPSs are attractive candidates for such co-products, with potential applications in cosmetics, pharmaceuticals and functional foods. Most SPS research on photosynthetic organisms has focused on large eukaryotic algae^34^. However, these production schemes rely on large-scale outdoor or mariculture-type cultivation, which is sensitive to environmental fluctuations and difficult to control genetically. In contrast, model cyanobacteria can be cultured under well-defined conditions, are amenable to genetic manipulation, and are supported by extensive genomic and omics resources, making them highly attractive candidate platforms for SPS production.

In bacterial metabolic engineering, achieving heterologous expression of complex metabolic pathways comprising many genes is an important objective for the production of valuable secondary metabolites. Although it is relatively straightforward to functionally express metabolic pathways catalyzed by soluble cytosolic enzymes, it is considerably more challenging to heterologously reconstruct complex membrane-associated systems such as EPS biosynthetic machinery. In bacteria, heterologous expression of exopolysaccharide biosynthetic systems has been achieved, but most reported studies have been conducted in *Escherichia coli* and *Lactococcus*, in which glycan biosynthesis has been most intensively investigated because of their medical relevance in pathogenicity and bioactivity^35,36^. In cyanobacteria, only in recent years have a few successful examples been reported in which large gene clusters, such as the nitrogen fixation *nif* cluster and the microcystin biosynthetic pathway, were functionally expressed in heterologous hosts^37,38^. However, these pathways are localized in the cytosol. To our knowledge, there has been no previous report in cyanobacteria of an entire large gene cluster encoding such a complex membrane-associated biosynthetic system being heterologously expressed and functionally reconstituted.

In this study, we achieved, to our knowledge, the first successful heterologous production of an SPS in cyanobacteria by rationally reconstituting and introducing the synechan biosynthesis and regulatory gene cluster *xss*, previously identified in the freshwater model cyanobacterium *S*.6803, into the non-SPS-producing strain *S.*7942. In other words, our primary contribution is to demonstrate that introducing a defined set of genes for SPS biosynthesis and its regulation is sufficient to induce substantial accumulation of extracellular SPS. As shown in the previous sections, the polysaccharide produced by the QRTP-A strain exhibited a degree of sulfation comparable to that of native synechan and a similar set of constituent monosaccharides, but differed in monosaccharide composition ratios and in the relative distribution between the RPS and CPS fractions. This difference represents a current limitation in that the native synechan structure has not yet been fully reconstructed; at the same time, it can also be interpreted as evidence that SPS composition is tunable by the host cellular context and by the design of the introduced *xss* gene set. Numerous structure-activity relationship studies on glycosaminoglycans and algal SPSs have shown that the physicochemical properties and biological functions of SPSs depend not only on their monosaccharide composition but also strongly on the degree and position of sulfation, that is, the sulfation pattern^34,39,40^. Therefore, although the SPS obtained in this work is best described as a “synechan-like” SPS rather than an exact copy of the original polymer, our results can also be viewed as an example of achieving synthetic-biology-based remodeling of SPS composition in cyanobacteria. Furthermore, our RNA-seq data provide concrete guidance for additional metabolic engineering aimed at further increasing SPS productivity and tuning polymer properties. In the QRTP-A strain, the strong repression of PEP synthase implies an overall decrease in carbon flux from PEP into the NDP-sugar biosynthetic network. In addition, downregulation of genes involved in sulfate uptake and sulfur assimilation in the SPS-producing state indicates that the cells entered a general stress-response mode in which the supply of PAPS, the sulfate donor for sulfotransferases, is likely diminished. Strengthening NDP-sugar supply and sulfur assimilation should therefore enable further increases in SPS production.

The synechan biosynthetic system Xss investigated in this study represents one example of a “cyanobacterial-type SPS biosynthetic unit”, comprising the transcriptional regulatory module *xssQ-xssR-xssS* together with Wzx/Wzy-dependent or ABC transporter-dependent EPS biosynthetic machineries and sulfotransferases. In cyanobacterial species that harbor homologues of XssQ and XssR and are predicted to possess synechan-like regulatory networks for SPS production, exemplified by *Aphanothece sacrum*, which produces the valuable SPS sacran, similar synthetic-biology approaches could in the future be used to manipulate endogenous SPS biosynthetic systems and to heterologously express them in other hosts. Furthermore, in heterotrophic bacteria such as *Escherichia coli*, several studies have already demonstrated the synthetic-biology-based production of chondroitin sulfate-like or heparan sulfate-like polysaccharides by combining bacterial EPS biosynthetic pathways with animal-derived sulfotransferases^4^. These precedents strongly suggest that, in cyanobacteria as well, diverse bacterial-type EPS biosynthetic machineries, including those originating from cyanobacteria, such as the synechan pathway, and animal- or bacterial-derived sulfotransferases could be combined to build photosynthesis-driven production platforms for a wide range of useful SPSs. Looking ahead, we envision photosynthesis-driven cyanobacterial SPS cell factories that use CO₂ as the sole or major carbon source to stably supply structurally diverse SPSs of both native and heterologous origin, with monosaccharide compositions and sulfation patterns precisely tailored to specific applications. Together, these considerations indicate that the present work provides a technical and conceptual foundation for synthetic-biology-driven SPS composition engineering in cyanobacteria.

## Materials and methods

### Culture conditions for cyanobacteria

The cyanobacterium *Synechococcus elongatus* PCC 7942 (*S*.7942) WT strain and its derivatives were grown photoautotrophically at 30°C under continuous white light illumination (30 μmol photons m^−2^·s^−1^) in BG-11 medium^41^ with 2% CO_2_ bubbling. When appropriate, spectinomycin (final concentration = 40 μg·mL^-1^) and chloramphenicol (final concentration = 10 μg·mL^-1^) were added to the media. Cell density was monitored at 750 nm. After pre-incubation in liquid BG-11 medium for several days, the cells were harvested and inoculated into fresh BG-11 medium at an optical density OD_750_ of 0.10. After 1 day of cultivation, IPTG was added to a final concentration of 0.1 mM, and the cultures were further incubated for an additional 5 days. For the growth curve, measurements were conducted in triplicate. Significance was determined using a paired *t*-test in Microsoft Excel. For observation of cell sedimentation or flotation, the cultures were subsequently incubated statically for an additional 2 days.

### Construction of plasmids and mutants

Primers used are listed in Table S1. Each DNA fragment was amplified using KOD One DNA polymerase (TOYOBO, Japan), PrimeSTAR MAX DNA polymerase and PrimeSTAR GXL DNA polymerase (Takara, Japan) and subcloned into the vector plasmids using In-Fusion HD Cloning Kit (TaKaRa, Japan). For construction of the *xssQ, xssR*, and *xssT* expression plasmid, these genes were first cloned into the pACYC vector while being linked by two synthetic RBSs (RBS1: 5’-ATCGATTCACTGTATAACATTAAGAAGGAGGATTACAAA-3’; RBS2: 5’-taaaaTAGTGGAGGttactag-3’). The vector backbone, the *xssQ* region, the *xssR* region, and the *xssT* region were PCR-amplified using the primer pairs pA-F/R, Q-FpA/RRBS1, R-FRBS1/RRBS2, and T-FRBS2/RpA, respectively. Using this construct as a template, the *xssQRT* region was then cloned into the pBR322 vector together with the upstream and downstream regions of the NS1 neutral site to generate the pBNS1-*xssQRT* construct. The vector backbone, NS1 upstream region, *xssQRT* region, and NS1 downstream region were PCR-amplified using the primer pairs pB-FNS1/RNS1, NS1up-F/R, QRT-FNS1/FNS2, and NS1down-F/R, respectively. For construction of the *xssP-A* expression plasmid, the *xssP-A* region was cloned into the previously constructed pYS1C-GFP plasmid^18^ to generate the pYS1C-*xssP-A* construct. The vector backbone and the *xssP-A* region were PCR-amplified using the primer pair pY-Fxss/Rxss for the vector backbone and the primer pairs xssPA-F1/R1 and xssPA-F2/R2 for the *xssP-A* region. The plasmids were introduced into *S*.7942 by natural transformation, and the resulting transformants were checked by PCR amplification using the primer sets (GC1-8) as shown in Fig.2a. The *S*.7942 wild-type strain transformed with pBNS1-*xssQRT* was designated *S*.7942 QRT, and the QRT strain transformed with pYS1C-*xssP-A* was designated *S*.7942 QRTP-A.

### Transcriptome analysis

Cells were cultured as in the culture conditions for cyanobacteria section and Fig. 2b and were collected on the third day (2 days after IPTG addition). Total RNA was extracted from the cells as previously reported^42^. Ribosomal RNA was subsequently removed from the total RNA using NEBNext rRNA Depletion Kit (Bacteria) (New England Biolabs). The sequencing libraries were generated according to the manufacturer’s instructions using NEBNext Ultra II Directional RNA Library Prep Kit for Illumina (New England Biolabs) with 200 bp library insertion size. The resulting 9 libraries were sequenced using the Illumina NextSeq 1000 sequencing platform. Three biological replicates were made for each condition. The reads were trimmed using CLC Genomics Workbench ver. 25.0.2. Trimmed reads were mapped to all genes in *S.*7942 (accession number: CP000100, CP000101) and the constructed plasmids (pBNS1-*xssQRT* and pYS1C-*xssP-A*) using CLC Genomics Workbench ver. 25.0.2 (QIAGEN, Hilden, Germany). The data underlying this article are available in the DRA/SRA database at [https://www.ddbj.nig.ac.jp/index-e.html] and can be accessed with the following accession numbers (DRR891100-DRR891108). The accession number of BioProject is PRJDB39866. Sequencing reads were mapped to the reference genome of *S.*7942. Gene expression levels were normalized and quantified as transcripts per million (TPM). Volcano plot analysis of the transcriptomes of the QRTP-A strain grown with or without IPTG was performed using iDEP (log2FC ≥ |1|, FDR-adjusted p-value ≤ 0.05)^43^.

### Scanning electron microscopy

SEM observations were performed by Hanaichi Ultrastructure Research Institute Co., Ltd. Cell samples for SEM were initially fixed in cacodylate-buffered 2% glutaraldehyde and 0.1% ruthenium red. Subsequently, they were post-fixed in 2% osmium tetroxide and 0.1% ruthenium red for 2 hours in an ice bath. Then, the specimens were dehydrated in a graded ethanol and dried by CO_2_ critical point dry. Dried specimens were coated by osmium plasma ion coater and were submitted to SEM observation (JSM-7500F at 5 kV).

### Alcian blue staining

The polysaccharides were stained with 1% Alcian blue 8GX (Merck) for 10 min in 3 % acetic acid (pH 2.5) or 0.5 N HCl (pH 0.5) as previously described ^44^.

### EPS fractionation

EPS fractionation was performed based on our previous method^6^. The entire culture was first centrifuged at 10,000 × g for 10 min to remove cells and CPS and then filtered through a 1.0-μ m pore PTFE membrane. The trapped materials were gently and carefully recovered from the membrane using MilliQ water with the aid of flat-tip tweezers. The sample collected was vortexed and then centrifuged at 20,000 × g for 10 min to remove cells. This fraction was used as the RPS sample. CPS was released from the cell pellet by vigorous vortexing with MilliQ water and recovered by centrifugation to remove cells (20,000 × g for 10 min).

### Sugar quantification

Total sugar was quantified using the phenol-sulfate method^45^. A 100-μL aliquot of 5% (w/w) phenol was added to 100 μL of a sample in a glass tube and vortexed three times for 10 s. Then, 500 μL of concentrated sulfuric acid was added, and the tube was immediately vortexed three times for 10 s and then kept at 30°C for 30 min in a water bath. Sugar content was measured by absorption at 487 nm using a UV-2600PC spectrophotometer (Shimadzu, Japan). Any contamination of the BG11 medium was evident by slight background coloration. This background was subtracted on the basis of the extrapolation of absorption at 430 nm, where the coloration due to sugars was minimal. Glucose was used as the standard. Some EPS samples were highly viscous, so we vortexed and sonicated them before measurement. Statistical significance was determined using Welch’s *t* test.

### Sugar composition analysis

Sugar composition analysis was performed based on our previous method^6^. The collected EPS samples were dialyzed with MilliQ water and then freeze-dried for 3 days. Sugar composition was analyzed by Toray Research Center, Inc. (Tokyo, Japan). A portion of the freeze-dried EPS sample was dissolved in 200 μL of 2 M trifluoroacetic acid and hydrolyzed at 100°C for 6 h. The treated sample was vacuum-dried with a centrifugal evaporator, redissolved in 400 μL MilliQ water, and filtered through a 0.22-μm pore filter. This sample was used for analysis.

Monosaccharide composition was determined by HPLC with the LC-20A system (Shimadzu). For neutral sugars, the column was TSK-gel Sugar AXG (TOSOH, Japan) and the temperature was 70°C. The mobile phase was 0.5 M potassium borate (pH 8.7) at 0.4 mL/min. Post-column labelling was performed using 1% (w/v) arginine and 3% (w/v) boric acid at 0.5 mL/min, 150°C. For uronic acids, the column was a Shimpack ISA-07 (Shimadzu) and the temperature was 70°C. The mobile phase was 1.0 M potassium borate (pH 8.7) at 0.8 mL/min. Post-column labelling was performed using 1% (w/v) arginine and 3% (w/v) boric acid at 0.8 mL/min, 150°C. The detector was a RF-10A_XL_ (Shimadzu), with excitation at 320 nm and emission at 430 nm. The standard curves were prepared for each monosaccharide with standard samples.

The SO_4_^2−^ content was determined by anion exchange column chromatography using the ISC-2100 system (Thermo Fisher Scientific, USA, Massachusetts). The column was eluted via a gradient of 0-1.0 M KOH. The separation column was IonPac ASI l-HC-4 μm (Thermo Fisher Scientific). Electrical conductivity was used for detection.

## Author information

### Authors

Kazuma Ohdate - *Department of Bioscience, Tokyo University of Agriculture, Tokyo 156-8502, Japan*

Yutaka Sakamaki - *Department of Bioscience, Tokyo University of Agriculture, Tokyo 156-8502, Japan*

Kaori Nimura-Matsune - *Department of Bioscience, Tokyo University of Agriculture, Tokyo 156-8502, Japan*

## Author contributions

K.M. and S.W. designed the concept and the experiments of this study; K.M., K.O., Y.S. and K.N-M. performed the experiments; K.M., K.O., K.N-M. and S.W. analyzed the data; K.M. and S.W. wrote the manuscript.

## Acknowledgment

We are grateful to Professor Emeritus Masahiko Ikeuchi for valuable comments on the concept of this study.

## Funding

This research was supported in part by JP19J01251 (K.M.), JST ACT-X grant no. JPMJAX20BG (K.M.) and Cooperative Research Grant of the Genome Research for BioResource from NODAI Genome Research Center, Tokyo University of Agriculture (K.M.).

### Abbreviations

EPS: exopolysaccharide;
SPS: sulfated polysaccharide;
CPS: capsular polysaccharides;
RPS: released polysaccharides;
NS: neutral site;
RBS: ribosome-binding site;
IPTG: isopropyl ß-D-1-thiogalactopyranoside;
WT: wild type;
Glc: D-glucose;
Gal: D-galactose;
Man: D-mannose;
Xyl: D-xylose.
WL: white light;
PEP: phosphoenolpyruvate.

## Data availability

The RNA-seq data generated in this study have been deposited in the DRA/SRA database at the DNA Data Bank of Japan under accession numbers DRR891100-DRR891108 (BioProject PRJDB39866). All other data supporting the findings of this study are available from the corresponding authors upon reasonable request.

## REFERENCES

1. Ragul, S., Sneha, S., Nautiyal, S., Balde, A. & Nazeer, R. A. Sulfated polysaccharides and their potential applications as drug carrier systems: A review. Reactive and Functional Polymers. 219, 106576 (2026).

2. Cimini, D., Bedini, E. & Schiraldi, C. Biotechnological advances in the synthesis of modified chondroitin towards novel biomedical applications. Biotechnology Advances. 67, 108185 (2023).

3. Rodriguez, M. L., Jann, B. & Jann, K. Structure and serological characteristics of the capsular K4 antigen of Escherichia coli O5: K4: H4, a fructose-containing polysaccharide with a chondroitin backbone. European journal of biochemistry. 177, 117–124 (1988).

4. Badri, A. et al. Complete biosynthesis of a sulfated chondroitin in Escherichia coli. Nature Communications. 12, 1389 (2021).

5. Kokoulin, M. S., Savicheva, Y. V., Filshtein, A. P., Romanenko, L. A. & Isaeva, M. P. Structure of a Sulfated Capsular Polysaccharide from the Marine Bacterium Cobetia marina KMM 1449 and a Genomic Insight into Its Biosynthesis. Mar. Drugs. 23, 29 (2025).

6. Maeda, K., Okuda, Y., Enomoto, G., Watanabe, S. & Ikeuchi, M. Biosynthesis of a sulfated exopolysaccharide, synechan, and bloom formation in the model cyanobacterium Synechocystis sp. strain PCC 6803. Elife. 10, e66538 (2021).

7. Flombaum, P. et al. Present and future global distributions of the marine Cyanobacteria Prochlorococcus and Synechococcus. Proc. Natl. Acad. Sci. 110, 9824–9829 (2013).

8. Mangan, N. M., Flamholz, A., Hood, R. D., Milo, R. & Savage, D. F. pH determines the energetic efficiency of the cyanobacterial CO2 concentrating mechanism. Proc. Natl. Acad. Sci. 113, E5354–E5362 (2016).

9. Angermayr, S. A., Rovira, A. G. & Hellingwerf, K. J. Metabolic engineering of cyanobacteria for the synthesis of commodity products. Trends in biotechnology. 33, 352–361 (2015).

10. Toepel, J., Karande, R., Klähn, S. & Bühler, B. Cyanobacteria as whole-cell factories: current status and future prospectives. Curr. Opin. Biotechnol. 80, 102892 (2023).

11. Pereira, S. et al. Complexity of cyanobacterial exopolysaccharides: composition, structures, inducing factors and putative genes involved in their biosynthesis and assembly. FEMS Microbiol. Rev. 33, 917–941 (2009).

12. Mouhim, R. F., Cornet, J.-F., Fontane, T., Fournet, B. & Dubertret, G. Production, isolation and preliminary characterization of the exopolysaccharide of the cyanobacterium *Spirulina platensis*. Biotechnol. Lett. 15, 567–572 (1993).

13. Okajima, M. K. et al. Supergiant ampholytic sugar chains with imbalanced charge ratio form saline ultra-absorbent hydrogels. Macromolecules. 41, 4061–4064 (2008).

14. Mota, R. et al. Cyanoflan: A cyanobacterial sulfated carbohydrate polymer with emulsifying properties. Carbohydrate Polymers. 229, 115525 (2020).

15. Ngatu, N. R. et al. Anti-inflammatory effects of sacran, a novel polysaccharide from Aphanothece sacrum, on 2, 4, 6-trinitrochlorobenzene–induced allergic dermatitis in vivo. Annals of Allergy, Asthma & Immunology. 108, 117–122. e112 (2012).

16. Maeda, K. et al. Genetic identification of factors for extracellular cellulose accumulation in the thermophilic cyanobacterium *Thermosynechococcus vulcanus*: proposal of a novel tripartite secretion system. Mol. Microbiol. 109, 121–134 (2018).

17. Kopf, M. et al. Comparative analysis of the primary transcriptome of *Synechocystis sp*. PCC 6803. DNA Res. 21, 527–539 (2014).

18. Sakamaki, Y., Maeda, K., Nimura-Matsune, K., Chibazakura, T. & Watanabe, S. Characterization of a cyanobacterial rep protein with broad-host range and its utilization for expression vectors. Front. Microbiol. 14, 1111979 (2023).

19. Pereira, S. B., Mota, R., Vieira, C. P., Vieira, J. & Tamagnini, P. Phylum-wide analysis of genes/proteins related to the last steps of assembly and export of extracellular polymeric substances (EPS) in cyanobacteria. Sci. Rep. 5, 14835 (2015).

20. Goclaw-Binder, H., et al. Nutrient-associated elongation and asymmetric division of the cyanobacterium Synechococcus PCC 7942. Environmental microbiology. 14, 680–690 (2012).

21. Kato, R., Maeda, K., Yano, T.-A., Tanaka, K. & Tanaka, T. Label-free visualization of photosynthetic microbial biofilms using mid-infrared photothermal and autofluorescence imaging. Analyst. 148, 6241–6247 (2023).

22. Goto-Seki, A., Shirokane, M., Masuda, S., Tanaka, K. & Takahashi, H. Specificity crosstalk among group 1 and group 2 sigma factors in the cyanobacterium Synechococcus sp. PCC7942: in vitro specificity and a phylogenetic analysis. Mol. Microbiol. 34, 473–484 (1999).

23. Suban, S. et al. A cyanobacterial sigma factor F controls biofilm-promoting genes through intra-and intercellular pathways. Biofilm. 8, 100217 (2024).

24. Flores, C. et al. The alternative sigma factor SigF is a key player in the control of secretion mechanisms in Synechocystis sp. PCC 6803. Environ Microbiol. 21, 343–359 (2019).

25. Maeda, K., Narikawa, R. & Ikeuchi, M. CugP is a novel ubiquitous non-GalU- type bacterial UDP-glucose pyrophosphorylase found in cyanobacteria. J. Bacteriol. 196, 2348–2354 (2014).

26. Suzuki, E. et al. Carbohydrate metabolism in mutants of the cyanobacterium Synechococcus elongatus PCC 7942 defective in glycogen synthesis. Appl. Environ. Microbiol. 76, 3153–3159 (2010).

27. Sára, M. & Sleytr, U. B. S-layer proteins. J. Bacteriol. 182, 859–868 (2000).

28. Flores, E. & Herrero, A. Nitrogen assimilation and nitrogen control in cyanobacteria (Portland Press Ltd., 2005).

29. Collier, J. L. & Grossman, A. A small polypeptide triggers complete degradation of light-harvesting phycobiliproteins in nutrient-deprived cyanobacteria. EMBO J. 13, 1039–1047 (1994).

30. Chen, H.-Y., Bandyopadhyay, A. & Pakrasi, H. B. Function, regulation and distribution of IsiA, a membrane-bound chlorophyll a-antenna protein in cyanobacteria. Photosynthetica. 56, 322–333 (2018).

31. Cruce, J. R. & Quinn, J. C. Economic viability of multiple algal biorefining pathways and the impact of public policies. Applied Energy. 233, 735–746 (2019).

32. Ubando, A. T., Ng, E. A. S., Chen, W.-H., Culaba, A. B. & Kwon, E. E. Life cycle assessment of microalgal biorefinery: A state-of-the-art review. Bioresource technology. 360, 127615 (2022).

33. Chew, K. W. et al. Microalgae biorefinery: high value products perspectives. Bioresource technology. 229, 53–62 (2017).

34. Jiao, G., Yu, G., Zhang, J. & Ewart, H. S. Chemical structures and bioactivities of sulfated polysaccharides from marine algae. Mar. Drugs. 9, 196–223 (2011).

35. Germond, J. E., Delley, M., D’Amico, N. & Vincent, S. J. Heterologous expression and characterization of the exopolysaccharide from Streptococcus thermophilus Sfi39. European Journal of Biochemistry. 268, 5149–5156 (2001).

36. Harding, C. M. et al. A platform for glycoengineering a polyvalent pneumococcal bioconjugate vaccine using E. coli as a host. Nature communications. 10, 891 (2019).

37. Tsujimoto, R. et al. Functional expression of an oxygen-labile nitrogenase in an oxygenic photosynthetic organism. Sci. Rep. 8, 7380 (2018).

38. Zheng, Y. et al. Reconstitution and expression of mcy gene cluster in the model cyanobacterium Synechococcus 7942 reveals a role of MC-LR in cell division. New Phytologist. 238, 1101–1114 (2023).

39. Pereira, M. S., Melo, F. R. & Mourão, P. A. Is there a correlation between structure and anticoagulant action of sulfated galactans and sulfated fucans? Glycobiology. 12, 573–580 (2002).

40. Shen, Q., Guo, Y., Wang, K., Zhang, C. & Ma, Y. A review of chondroitin sulfate’s preparation, properties, functions, and applications. Molecules. 28, 7093 (2023).

41. Stanier, R., Kunisawa, R., Mandel, M. & Cohen-Bazire, G. Purification and properties of unicellular blue-green algae (order *Chroococcales*). Bacteriol. Rev. 35, 171–205 (1971).

42. Watanabe, S. et al. Regulation of RNase E during the UV stress response in the cyanobacterium Synechocystis sp. PCC 6803. Mlife. 2, 43–57 (2023).

43. Ge, S. X., Son, E. W. & Yao, R. iDEP: an integrated web application for differential expression and pathway analysis of RNA-Seq data. BMC bioinformatics. 19, 534 (2018).

44. Di Pippo, F. et al. Characterization of exopolysaccharides produced by seven biofilm-forming cyanobacterial strains for biotechnological applications. J. Appl. Phycol. 25, 1697–1708 (2013).

45. DuBois, M., Gilles, K. A., Hamilton, J. K., Rebers, P. T. & Smith, F. Colorimetric method for determination of sugars and related substances. Anal. Chem. 28, 350–356 (1956).

