## Supplementary Information for "Transfer of the synechan biosynthesis and regulatory pathway enables sulfated polysaccharide production in *Synechococcus elongatus* PCC 7942"

*** Correspondence:**
Co-corresponding Author

### 1. Data

**Supplementary Data S1: Sequence of plasmid pYS1C-*xssP-A*.**

**Supplementary Data S2: Sequence of plasmid pBNS1-*xssQRT*.**

### 2. Figures

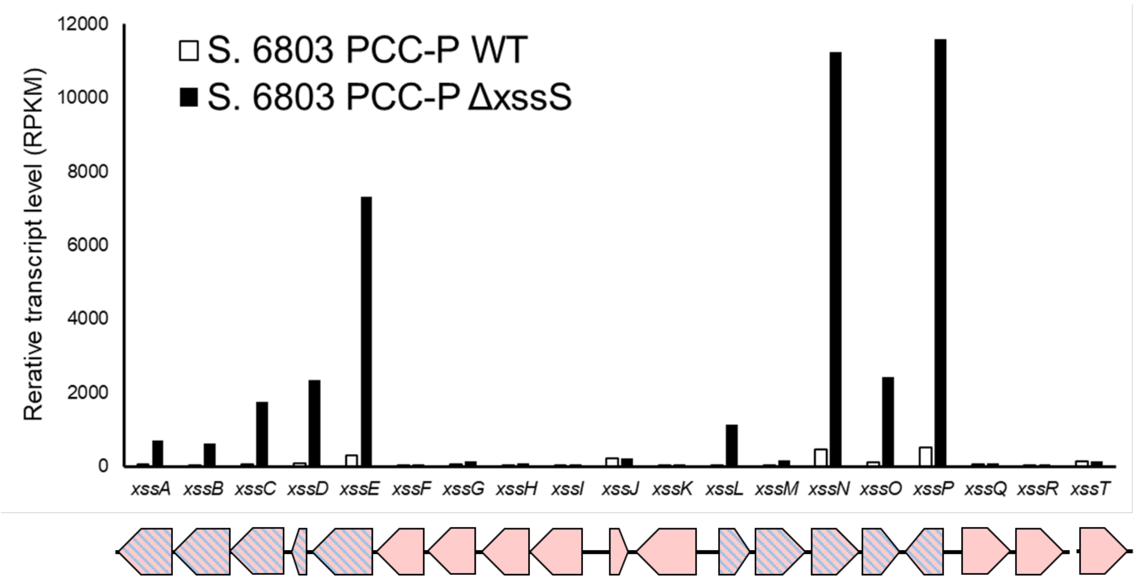


**Figure S1. Gene expression analysis of the *xss* gene cluster based on RNA-seq in our previous research[1].**

White bars and black bars represent transcript levels of *S.*6803 PCC-P WT and Δ*xssS* mutant, respectively. The schematic below the graph shows the arrangement of the *xss* genes corresponding to the bars in the graph. Each pentagon represents a gene; pentagons with pink and blue stripes indicate genes that are under XssQ-dependent transcriptional regulation in *S.*6803, and pink pentagons indicate the remaining *xss* genes.

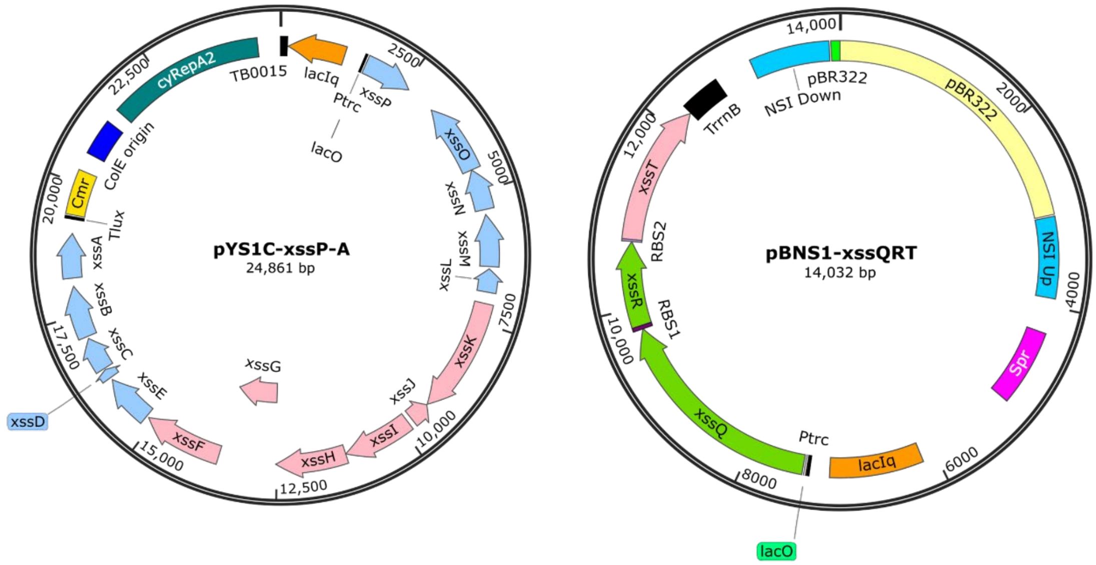


**Figure S2. Maps of the plasmids used in this study**

Plasmid maps of pYS1C-*xssP-A* and pBNS1-*xssQRT*, used in this study to construct the heterologous expression strain for the synechan biosynthetic pathway.

These maps were created with SnapGene (reference).

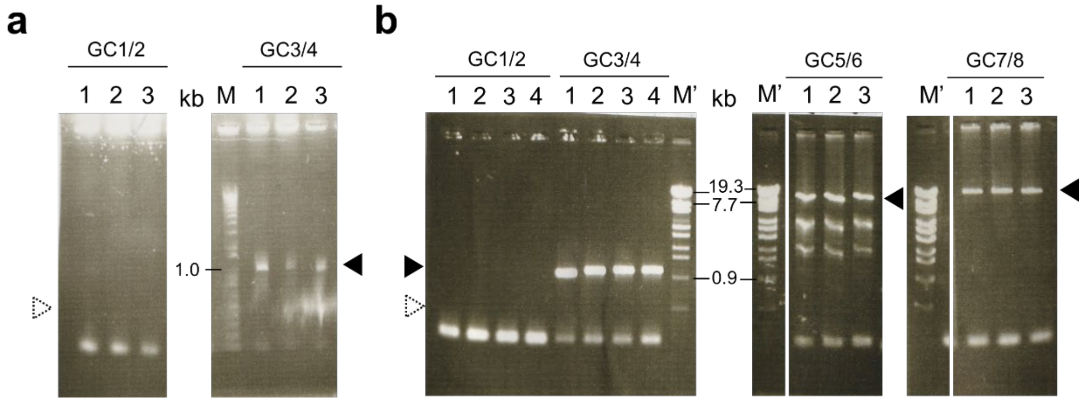


**Figure S3. Agarose gel electrophoresis of PCR products to assess gene integration and segregation of mutants.**

Genomic DNA extracted from the two recombinant strains constructed in this study, (a) QRT and (b) QRTP-A, was used as a template for PCR with the primer sets shown in Fig. 2a, and the resulting products were analyzed by agarose gel electrophoresis. M, 1 kb ladder marker; M', λ-EcoT14I digest marker. The numbers above each lane indicate individual clones. The expected sizes of the PCR products for each primer set were as follows: 1/2, 403 bp; 3/4, 891 bp; 5/6, 8417 bp; and 7/8, 9238 bp. The presence or absence of bands is indicated by filled and open arrowheads, respectively. The two images in (a) and the four images on the right in (b) were each derived from the same gel, but for clarity, unnecessary lanes were removed, and the removed regions are indicated by white lines.

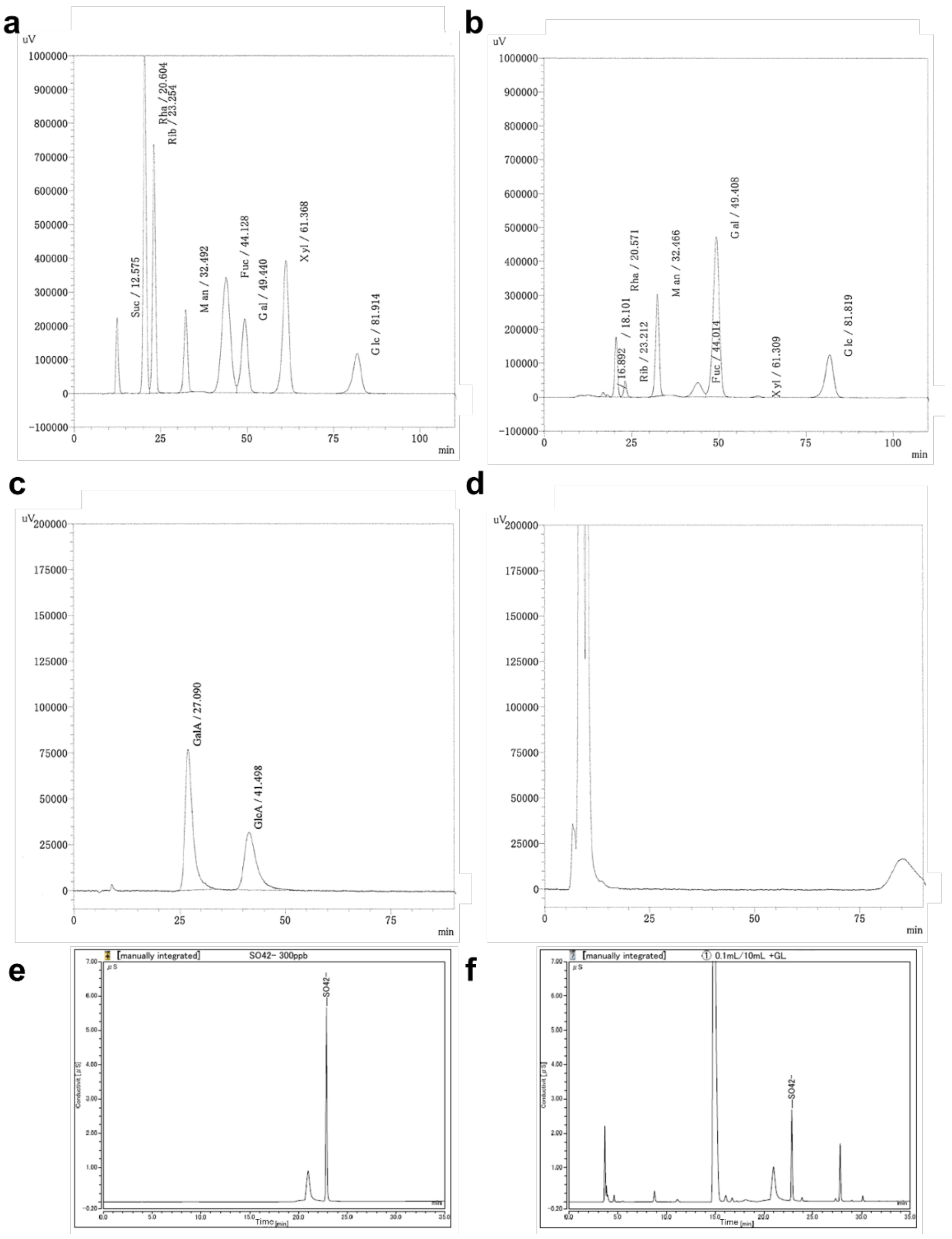


**Figure S4.** Chromatograms of HPLC and anion exchange column chromatography of the *S.*7942 QRTP-A RPS.

**a** and **b**, HPLC profiles for neutral sugars of standards (a) and RPS from *S.*7942 QRTP-A mutant strain (b). **c** and **d**, HPLC profiles for uronic acids of standards (c) and RPS from *S.*7942 QRTP-A mutant strain (d). The corresponding monosaccharide and retention time are noted at each peak. **e** and **f**, HPLC profiles for $SO_4^{2-}$ after hydrolysis of standards (e) and RPS from *S.*7942 QRTP-A mutant strain (f).

### 3. Table

**Table S1. Oligonucleotide primers used in this study.**

| Purpose | Name | Sequence (5' => 3') |
|---|---|---|
| Strain construction | pA-F | TCCTCTACGCCGGACGCAT |
| | pA-R | CATGGTTTATTCCTCCTTATTT |
| | Q-FpA | GAGGAATAAACCATGGCAAAACGTTCCCTTAAAG |
| | Q-RRBS1 | CTTCTTAATGTTATACAGTGAATCGATTTAGCTAACCTTTTCGCC |
| | R-FRBS1 | TATAACATTAAGAAGGAGGATTACAAAATGACTGAGACTTCTCCG |
| | R-RRBS2 | GTAACCTCCACTATTTTACTACCCCATGACCAGGGC |
| | T-FRBS2 | AATAGTGGAGGTTACTAGATGAATGCTATGAATCCT |
| | T-RpA | GTCCGGCGTAGAGGATTAATTAAGGCGGTCGAT |
| | pB-FNS1 | AGAAAGCAGCGCTCCTGGTCATAGCTGTTTCCTGC |
| | pB-RNS1 | ATGCGGAGCGCTTTTTCACTGGCCGTCGTTTTACC |
| | NS1up-F | AAAAGCGCTCCGCATGGATCTGACC |
| | NS1up-R | GAATTCCATGGTCTGTTTCCTGTGT |
| | QRT-FNS1 | ACAGACCATGGAATTCGCAAAACGTTCCCTTAAAGC |
| | QRT-RNS1 | CAAAACAGCCAAGCTTTTAATTAAGGCGGTCGATGA |
| | NS1down-F | AAGCTTGGCTGTTTTGGCGGATGAG |
| | NS1doen-R | GGAGCGCTGCTTTCTTGGCAAGCGG |
| | pY-Fxss | AGAGTATCAAACCCCTAAGCTTACTAGTAATACTGCAGAG |
| | pY-Rxss | TTGAATAGTGCTAGCCATGATAACCTCCTAAATTGTTATCCGC |
| | xssPA-F1 | GCTAGCACTATTCAACTGATTGAAGTTCCCCAAAG |
| | xssPA-R1 | TGGGTCGAACAGAAATACGCTCTACGACAACACCA |
| | xssPA-F2 | GGGGTTTGATACTCTTCCAGCAACTCGTCTAGTAA |
| | xssPA-R2 | TTTCTGTTCGACCCATTGCCCAGACTGATTTAGCA |
| Genotype check | GC1 | CAAACAGGTGCAGCAGCAACT |
| | GC2 | CATCGCTATCTCTTAGGACTTCGCAG |
| | GC3 | GTCACCCTAAGAGATGGT |
| | GC4 | CAAAACAGCCAAGCTTTTAATTAAGGCGGTCGATGA |
| | GC5 | GCTAGCACTATTCAACTGATTGAAGTTCCCCAAAG |
| | GC6 | TGGGTCGAACAGAAATACGCTCTACGACAACACCA |
| | GC7 | TTTCTGTTCGACCCATTGCCCAGACTGATTTAGCA |
| | GC8 | GGGGTTTGATACTCTTCCAGCAACTCGTCTAGTAA |

**Table S2. Source data for RNA-seq analysis.** (additional Excel file)

**Table S3. Genes significantly upregulated upon IPTG induction in the QRTP-A strain.** (additional Excel file)

Genes listed in Table 2 are highlighted with a yellow background.

**Table S4. Genes significantly downregulated upon IPTG induction in the QRTP-A strain.** (additional Excel file)

Genes listed in Table 3 are highlighted with a yellow background.

### <u>Reference</u>

1. Maeda, K., Okuda, Y., Enomoto, G., Watanabe, S. & Ikeuchi, M. Biosynthesis of a sulfated exopolysaccharide, synechan, and bloom formation in the model cyanobacterium Synechocystis sp. strain PCC 6803. *Elife*. **10**, e66538 (2021).
